# The m^6^A writer FIONA1 methylates the 3’UTR of *FLC* and controls flowering in Arabidopsis

**DOI:** 10.1101/2022.01.24.477497

**Authors:** Bin Sun, Kaushal Kumar Bhati, Ashleigh Edwards, Louise Petri, Valdeko Kruusvee, Anko Blaakmeer, Ulla Dolde, Vandasue Rodrigues, Daniel Straub, Stephan Wenkel

**Author notes:** Correspondence: Stephan Wenkel.

## Abstract

Adenosine bases of RNA can be transiently modified by the deposition of a methyl-group to form N^6^-methyladenosine (m^6^A). This adenosine-methylation is an ancient process and the enzymes involved are evolutionary highly conserved. A genetic screen designed to identified suppressors of late flowering transgenic Arabidopsis plants overexpressing the miP1a microProtein yielded a new allele of the FIONA1 (FIO1) m^6^A-methyltransferase. To characterize the early flowering phenotype of *fio1* mutant plants we employed an integrative approach of mRNA-seq, Nanopore direct RNA-sequencing and meRIP-seq to identify differentially expressed transcripts as well as differentially methylated mRNAs. We provide evidence that FIO1 is the elusive methylase responsible for the 3’-end methylation of the *FLOWERING LOCUS C* (*FLC*) transcript. Furthermore, our genetic and biochemical data suggest that 3’-methylation stabilizes *FLC* mRNAs and non-methylated *FLC* is a target for rapid degradation.

## INTRODUCTION

Modification of RNA is pervasive and found across the entire tree of life (Zaccara *et al*, 2019). Most abundant is the reversible conversion of adenine bases to N6-methyladenosine (m^6^A) in mRNA. In plants, m^6^A methylation patterns have been found to be highly conserved between distant ecotypes (Luo *et al*, 2014) suggesting ancient functions. In addition, loss of the METTL3-related methyltransferase MTA causes arrested development (Zhong *et al*, 2008), implying that m^6^A-methylation is both abundant and essential. Biochemical studies have revealed that the m^6^A-writer complex consists of METTL3, METTL14, and associated proteins (Liu *et al*, 2014). Besides METTL3 and METTL14, the human METTL16 methylase is also implicated in controlling m^6^A-methylation of mRNAs and snRNA (Pendleton *et al*, 2017) and has been shown in worms to affect diet-induced splicing of mRNA transcripts (Mendel *et al*, 2021). In plants, the functions of METTL3 (MTA) and METTL14 (MTB) (Růžička *et al*, 2017) as m^6^A-methylation writers are well characterized. In addition to m^6^A-writers, m^6^A-reader complexes can recognize m^6^A marks and affect RNA stability, splicing and translation (Arribas-Hernández *et al*, 2018). The analysis of an early flowering knock-down allele of the METTL16-homolog FIONA1, *fio1-2*, revealed changes in the m^6^A methylation status of many genes, several encoding flowering regulators including *SUPPRESSOR OF OVEREXPRESSION OF CONSTANS* (*SOC1*) (Xu *et al*, 2022). Besides *SOC1* mRNA, the mRNA of the flowering regulator *FLOWERING LOCUS C* (*FLC*) has also been shown to be modified by m^6^A-methylation (Xu *et al*, 2021). The latter study showed that an R-loop forms at the FLC locus that is resolved by the RNA-binding proteins FCA and FY. In this process, FCA binds the *FLC COOLAIR* antisense transcript to facilitate m^6^A-methylation (Xu *et al*., 2021). Interestingly, the authors also detected m^6^A-methylation of the 3’UTR of *FLC* mRNA but this methylation appeared to be FCA-independent.

Here, we isolated a novel allele of *FIONA1* (*FIO1*) in a genetic screen for suppressors of the late flowering phenotype of plants overexpressing the miP1a microProtein (Graeff *et al*, 2016). We present evidence that FIO1 acts as m^6^A-methyltransferase in Arabidopsis and is the functional homolog of the human METTL16. Using a combination of mRNA-seq, meRIP-seq and Nanopore direct RNA-sequencing, we provide further evidence that FIO1 is the elusive 3’UTR methylase of *FLC*. Moreover, our data shows that the largely pleiotropic phenotype of *fio1* mutant plants is a result of massive transcriptome and RNA-methylome changes. In the case of *FLC*, FIO1 is needed to maintain the 3’-end methylation. Abrogation of this methylation mark causes depletion of *FLC* mRNA.

## RESULTS

### FIONA1 acts as a floral repressor that functions partially independent of the photoperiod pathway

The miP1a/miP1b microProteins act as suppressors of flowering by interacting with a TOPLESS-containing repressor complex (Graeff *et al*., 2016; Rodrigues *et al*, 2021). To identify factors that are required for the repressor complex to suppress flowering, we performed a genetic screen with transgenic *miP1a-OX* (*35S::MIP1A*) plants. We identified a set of *<u>su</u>ppressor of <u>mi</u>P1a* (*sum*) mutants, that, despite high levels of miP1a protein, flowered early under inductive long day conditions (Rodrigues *et al*., 2021). One of the suppressors, *sum8*, we describe here, showed accelerated flowering compared to the non-mutagenized *miP1a-OX* parental plant (Fig. 1a,b). To identify the causal mutation in the *sum8* background, we crossed *miP1a-OX sum8* plants to Col-0 wildtype, self-pollinated the offspring and selected a pool of 20 BASTA-resistant suppressor mutants of the following generation. Pooled DNA of the *sum8* suppressor mutant and the parental line was then analyzed by genome re-sequencing. In total, we detected 685 EMS-induced SNPs with a frequency enrichment in the middle of chromosome 2 (Fig. 1c). At the summit region of the enrichment peak we identified a point mutation in the *FIONA1* (*FIO1*) gene which converted the serine 278 into an asparagine (S278N). To verify that the mutation in *FIO1* is causal for the early flowering phenotype, we obtained a second EMS allele (*fio1-1*) that had been described earlier (Kim *et al*, 2008) and crossed it with *miP1a-OX sum8* plants. The resultant nullizygote offspring (*miP1a-OX/+ fio1-1/sum8*) flowered early (Fig. 2), supporting that the mutation in *FIO1* is indeed causal for the flowering phenotype. The *fio1-1* allele is a splice site mutation that results in the loss of five amino acids while *sum8* is a point mutation. To obtain an additional *FIO1* allele, we used a CRISPR approach with multiple sgRNAs and obtained the new allele *fio1-3*. Like *fio1-1* and *sum8*, also *fio1-3* showed early flowering in long day conditions (Supplementary figure 1a, b). The *fio1-3* deletion occurred close to a splice site and caused the loss of amino acids 53-64 and the amino acid conversions of residues 66-72 (Supplementary Figure 1c).

**Figure 1.**
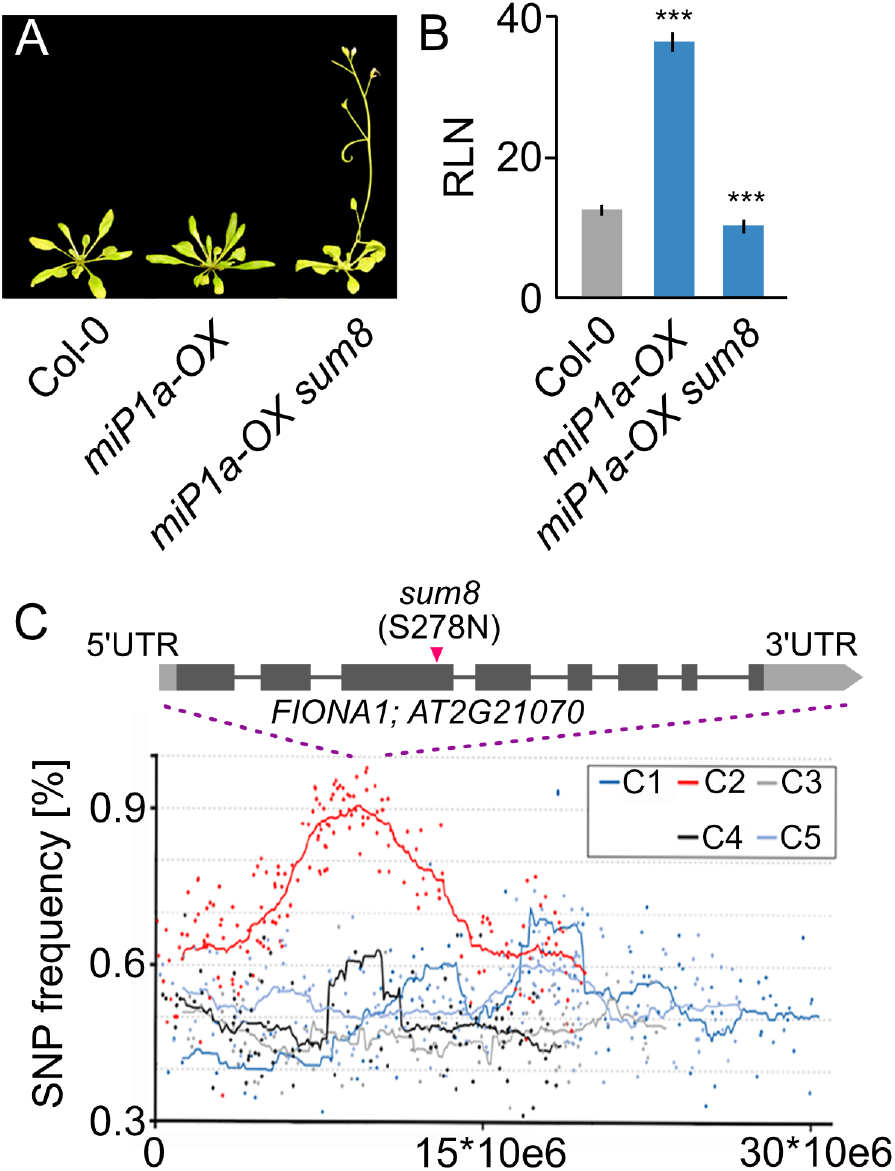
Identification of flowering repressor FIONA1 by whole-genome re-sequencing. **(A)** Phenotype of the *sum8* (*fio1*) mutant in the miP1a-OX background compared to the Col-0 wildtype grown in LD conditions. **(B)** Determination of flowering by counting the number of rosette leaves (RLN = rosette leaf number) at the bolting stage in LD. Plotted are average leaf number +/- SD, ***p=<0.001, N=10. **(C)** Mapping-by-sequencing of the *sum8* suppressor mutation. Plotted are SNP frequencies of a pool of segregating F2 plants. Increased SNP frequencies were observed in chromosome 2 and the FIO1 locus is at the summit of the plot.

**Figure 2.**
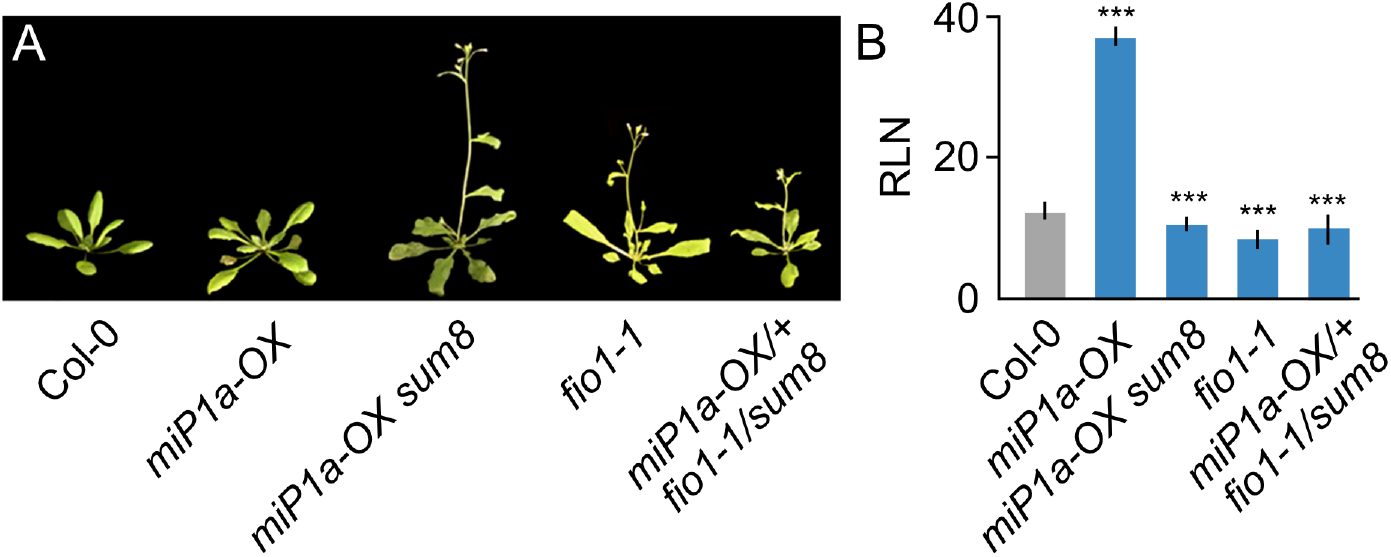
FIO1 is the gene affected by the *sum8* mutation. **(A)** Genetic complementation experiment proving that the sum8 mutation affects FIO1. Shown are the flowering phenotypes of plants grown in LD conditions. **(B)** Determination of flowering by counting the number of rosette leaves (RLN = rosette leaf number) at the bolting stage in LD. Plotted are average leaf number +/- SD, ***p=<0.001, N=10.

### The loss of FIO1 function affects multiple flowering pathways

A previous genetic screen for regulators of flowering resulted in the identification of the *fio1-1* mutant that exhibited early flowering in both long- and short-day conditions (Kim *et al*., 2008). A knock-down mutation caused by a T-DNA insertion in the 5’-region of the *FIONA1* gene (Xu *et al*., 2022) showed a similar phenotype. The *fio1-1* mutant was shown to have elevated levels of both *CONSTANS* (*CO*) and *FLOWERING LOCUS T* (*FT*) mRNA. CO is a photoperiod-sensitive transcription factor that accumulates in response to long days to activate *FT* (Valverde *et al*, 2004), which in turn acts as florigen to induce flowering (Corbesier *et al*, 2007; Tamaki *et al*, 2007). The flowering phenotype of *fio1-1* was ascribed to changes in period length of the central oscillator. Consistent with previous findings, we found that levels of both *CO* and *FT* were elevated in *fio1-1* and *fio1-3* (Fig. 3a,b). A genetic interaction study revealed that *miP1a miP1b fio1-3* triple mutant plants flowered early like *fio1-3* mutant plants. The combination of *fio1* mutants with either *co* and *ft* mutants as in *fio1 co* and *fio1 ft*, revealed a promotion of flowering (Fig. 3c,d) in both short days and long days. These results unequivocally show that the function of FIO1 is independent of the function of miP1a.

**Figure 3.**
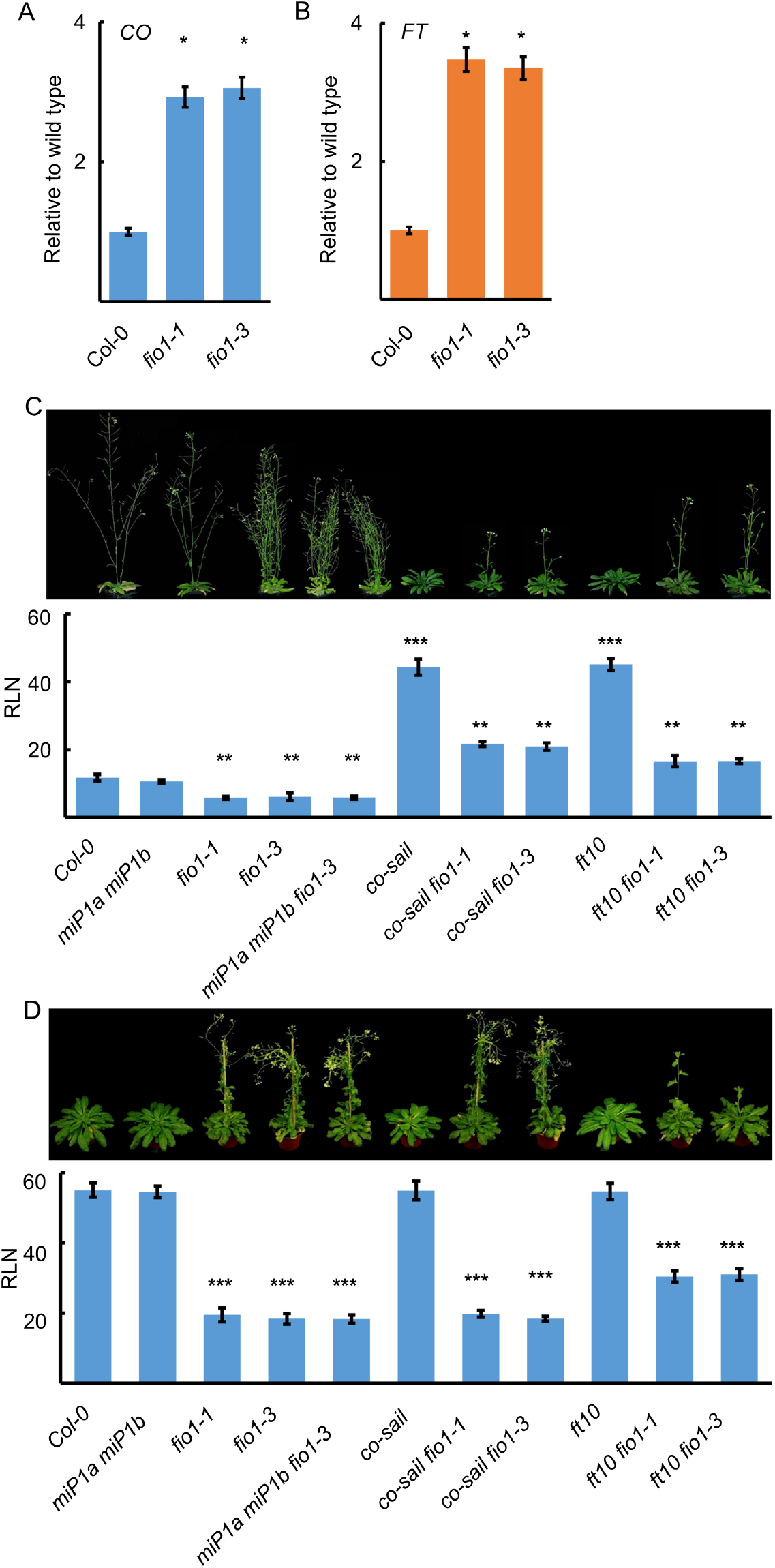
FIONA1 acts partially independent of the photoperiod pathway to repress flowering. **(A)** and **(B)** Quantification of *CO* and *FT* in Col-0, *fio1-1* and *fio1-3* by qRT-PCR. Values are the means ±SD. N = 4. * P ≤ 0.01. **(C)** Phenotypes of *miP1a miP1b, fio1-1, fio1-3, miP1a miP1b fio1-3, co-sail, co-sail fio1-1, co-sail fio1-3, ft10, ft10 fio1-1, ft10 fio1-3* and determination of flowering time by counting the number of rosette leaves at bolting compare to wild type, under long day conditions. RLN = number of rosette leaves at the bolting stage. Values are the means ±SD. N = 10 to 20. One-way ANOVA was carried out to test significance, **P ≤ 0.005, ***P≤ 0.001. **(D)** Phenotypes of *miP1a miP1b, fio1-1, fio1-3, miP1a miP1b fio1-3, co-sail, co-sail fio1-1, co-sail fio1-3, ft10, ft10 fio1-1, ft10 fio1-3* and determination of flowering time by counting the number of rosette leaves at bolting compare to wild type, under short day conditions. RLN = number of rosette leaves at the bolting stage. Values are the means ±SD. N = 10 to 12. One-way ANOVA was carried out to test significance, ***P≤ 0.001.

### Transcriptome analysis of *fio1-1* and *fio1-3* mutant plants

To obtain a better understanding of how FIO1 affects flowering, we performed an RNA-seq experiment with Col-0, *fio1-1* and *fio1-3* mutant plants to identify differentially expressed genes. RNA of two biological replicates of 14 day-old seedlings was isolated and sequenced on an Illumina HiSeq instrument. After removing low-quality reads, an average of 91.47% of the filtered reads was mapped to the *Arabidopsis thaliana* reference genome. Principal component analysis (PCA) and hierarchical cluster analysis (HCA) revealed that the individual biological replicates clustered closely together (Fig. 4a, b), indicating a high degree of experimental reproducibility. Interestingly, *fio1-1* and *fio1-3* were also distinct from each other and wild type, indicating that although they show a similar flowering phenotype they might differ at the molecular level.

**Figure 4.**
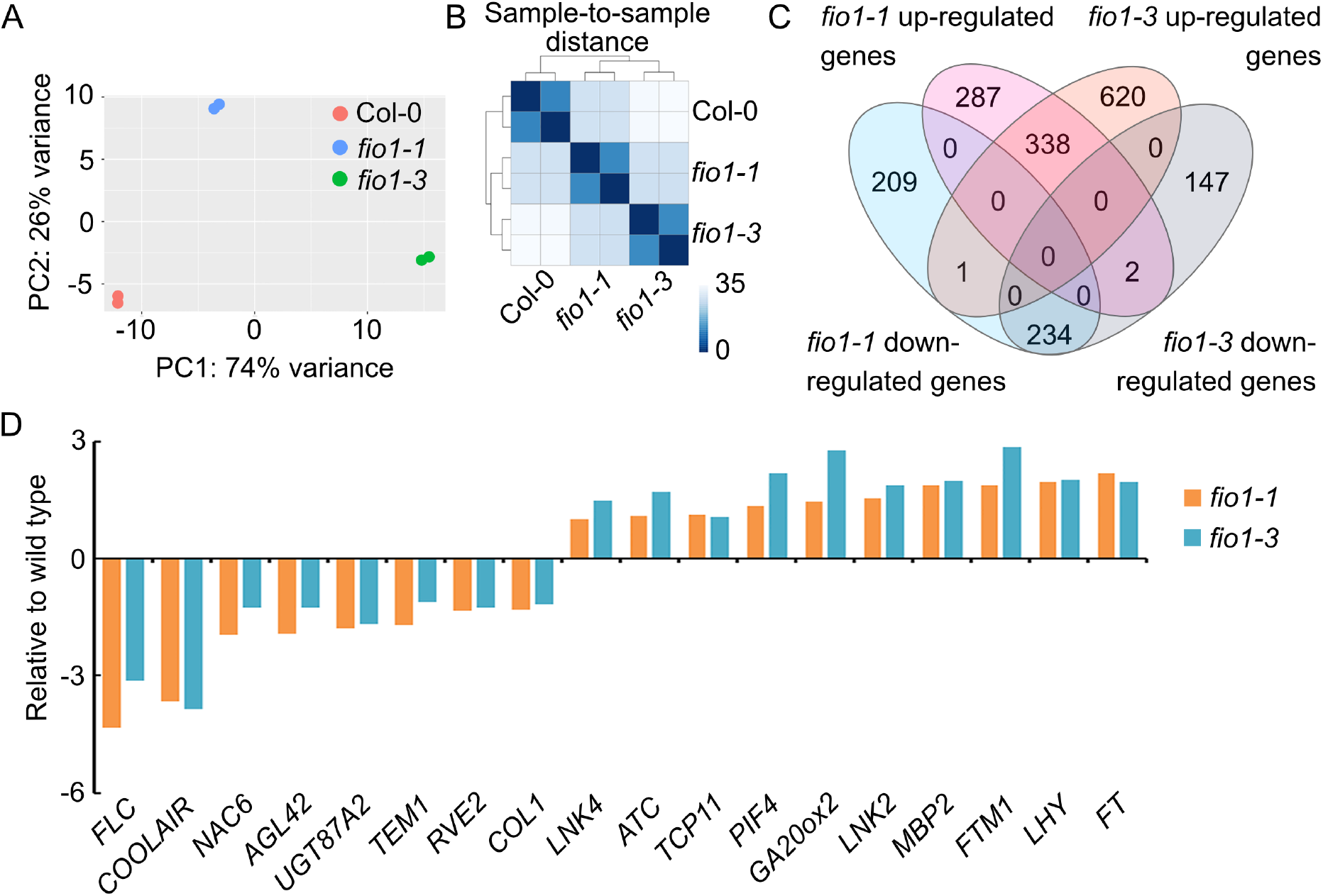
Transcriptome changes observed in *fio1* mutants. **(A)** Principal component analysis (PCA) plot displaying the different RNA-seq performed using DESeq2 rlog-normalized RNA-seq data. Plotted is the percentage of variance for each component. **(B)** Hierarchical clustering analysis (HCA) of the different RNA-seq libraries. The heatmap was built using the DEseq2 package. Samples were clustered using HCA performed with DESeq2 rlog-normalized RNA-seq data, and the dendrogram represents the clustering results. The heatmap illustrates the pairwise distances between the different samples, with higher similarity indicated by higher intensity of color. **(C)** Venn diagram showing the overlap of differentially expressed genes in *fio1-1* and *fio1-3* compared to the wild type. The absolute value of log2 FC (fold change; *fio1* mutant / WT) ≥ 1.0 and adjusted P-value (false discovery rate; FDR) ≤ 0.05. **(D)** RNA-seq showing the expression levels of flowering related genes in *fio1-1* and *fio1-3* compared to the wild type. The absolute value of log2 FC (fold change; *fio1* mutant / WT) ≥ 1.0 and adjusted P-value (false discovery rate; FDR) ≤ 0.05.

To identify differentially expressed genes (DEGs) in *fio1-1* and *fio1-3* we used limma-voom (Law *et al*, 2014) with a fold change cutoff of 2.0 or more. In total, we identified 627 and 959 up-regulated genes in *fio1-1* plants and *fio1-3* plants respectively (P value < 0.05 and adjusted P value < 0.05; Supplementary Table 1). In total we found 1071 DEGs in fio1-1 and 1342 DEGs in fio1-3 with an overlap of 338 up-regulated genes and 234 down-regulated genes (Fig. 4c). A total of 18 misregulated genes were associated with regulation of flowering (Fig. 4d), these include the flowering repressors *FLOWERING LOCUS C* (*FLC*) and *TEMPRANILLO1* (*TEM1*) whose mRNA levels were significantly reduced in *fio1* mutant plants and the flowering activators *PHYTOCHROME INTERACTING FACTOR4* (*PIF4*), *FT* and *LATE ELONGATED HYPOCOTYL* (*LHY*) whose mRNA levels were significantly increased in *fio1* mutant plants (Fig. 4d). These findings are in agreement with the early flowering phenotype of *fio1* mutant plants.

### FIO1 is related to the human METTL16 protein

FIO1 is a nuclear localized protein containing a DUF890 domain that part of METTL16-like protein family comprising among others the human and mouse METTL16 and the C. elegans METT-10 proteins. Animals carrying loss-of-function alleles of METT-10/METTL16 have been described to show severe developmental defects, and sometimes, lethality (Dorsett *et al*, 2009; Mendel *et al*, 2018). The mutant phenotypes we observed in plants were rather mild regarding overall plant morphology which raised the question whether we were dealing with loss-of-function or reduced function alleles of *FIO1*. All mutants had either smaller deletions or a single amino acid change suggesting they could be weak, reduced function alleles. To gain further insights into the alleles that we had obtained, we created a homology model of the FIO1 methyltransferase (MTase) domain and compared it against the crystal structure of the human homologue, METTL16. In the case of the *sum8* mutation (S278N, Supplementary Figure 2), we found that the sidechain of S278 normally forms hydrogen bonds with the nitrogen on the W330 within the protein core. Upon mutating the serine to an asparagine, we expect that the larger asparagine sidechain cannot be accommodated in the protein interior, leading to disrupted domain fold and function. The *fio1-1* mutation involves the deletion of five amino acids 145-149 in the FIO1 protein (Supplementary Figure 2) which includes the disruption of a potential hydrogen bond between the sidechains of Q82 and T147 and the loss of a flexible loop connecting an alpha helix and a beta sheet. The *fio1-3* mutation involves the large deletion of amino acids 57-68 and the non-conservative mutation of residues 53-56 and 69-72 (Supplementary Figure 2). Both *fio1-1* and *fio1-3* involve the large-scale disruption of hydrophobic and hydrogen bonding interactions and are likely to result in misfolded or aggregated protein. Thus, it is highly likely that all three mutations (*sum8, fio1-1* and *fio1-3*) disrupt the methyltransferase function of FIO1.

To validate the findings of the protein modeling we employed a second CRISPR mutagenesis approach and designed eight sgRNAs spanning the entire *FIO1* locus and transformed these in bulk to obtain larger structural mutations (Supplementary Fig. S3). We identified 11 new FIO1 alleles several of which had large structural deletions. Three new alleles (*fio1-cr4, fio1-cr9, fio1-cr10*) had frame-shift mutations that would not lead to the production of functional proteins. All new alleles were viable and, apart from early flowering did not show severe developmental defects. Taken together, these results show that the loss of METTL16 function is not lethal in plants but affects the transition to flowering.

### FIO1 acts as m^6^A-methylase and methylates predominantly the 3’UTR of mRNAs

The presence of the DUF890 domain suggests that FIO1 acts as a genuine m^6^A methylase. To identify the FIO1 RNA substrates, we employed a modified version of methylated RNA-immunoprecipitation (meRIP) followed by deep sequencing that was described earlier (Fig. 5a) (Dominissini *et al*, 2013). To determine methylation positions (m^6^A peaks) we used MACS (Zhang *et al*, 2008) with a false discovery rate (FDR) ≤ 0.05 and enrichment of ≥ 2-fold of sequence reads. In summary, we identified 2,822, 2,375 and 2,580 m6A-methylation peaks in wild type, *fio1-1* and *fio1-3*, respectively (Supplementary Table. 2). In *fio1-1* plants and *fio1-3* plants we identified 80 and 143 peaks respectively with increased m^6^A level compared to wild type. In contrast, a total of 850 m^6^A methylation peaks in *fio1-1* and 989 peaks in *fio1-3* were decreased or absent compared to the wild type (Fig. 5b). These findings suggest that FIO1 methylates mRNAs. When assessing the localization of the m^6^A-peaks globally in wild type, *fio1-1* and *fio1-3*, we observed more peaks in exons of *fio1* mutants and a reduced number of peaks in the 3’UTR of *fio1* mutants compared to wild type (Fig. 5c). The differential m^6^A peak distribution analysis (wild type versus *fio1* mutants) revealed a massive over-representation of hypomethylated peaks in 3’UTRs in *fio1* mutants compared to wild type (Fig. 5d). The findings indicate that FIO1 acts as m^6^A methylase and methylates predominantly the 3’UTRs of its target substrates. To explore a potential connection between m^6^A-methylation and RNA stability we compared our mRNA-seq and MeRIP datasets. In total we found nine genes containing hypomethylated peaks, eight of which were expressed at lower levels while one was expressed at higher level in *fio1* mutants compared to the wild type (Fig. 5e).

**Figure 5.**
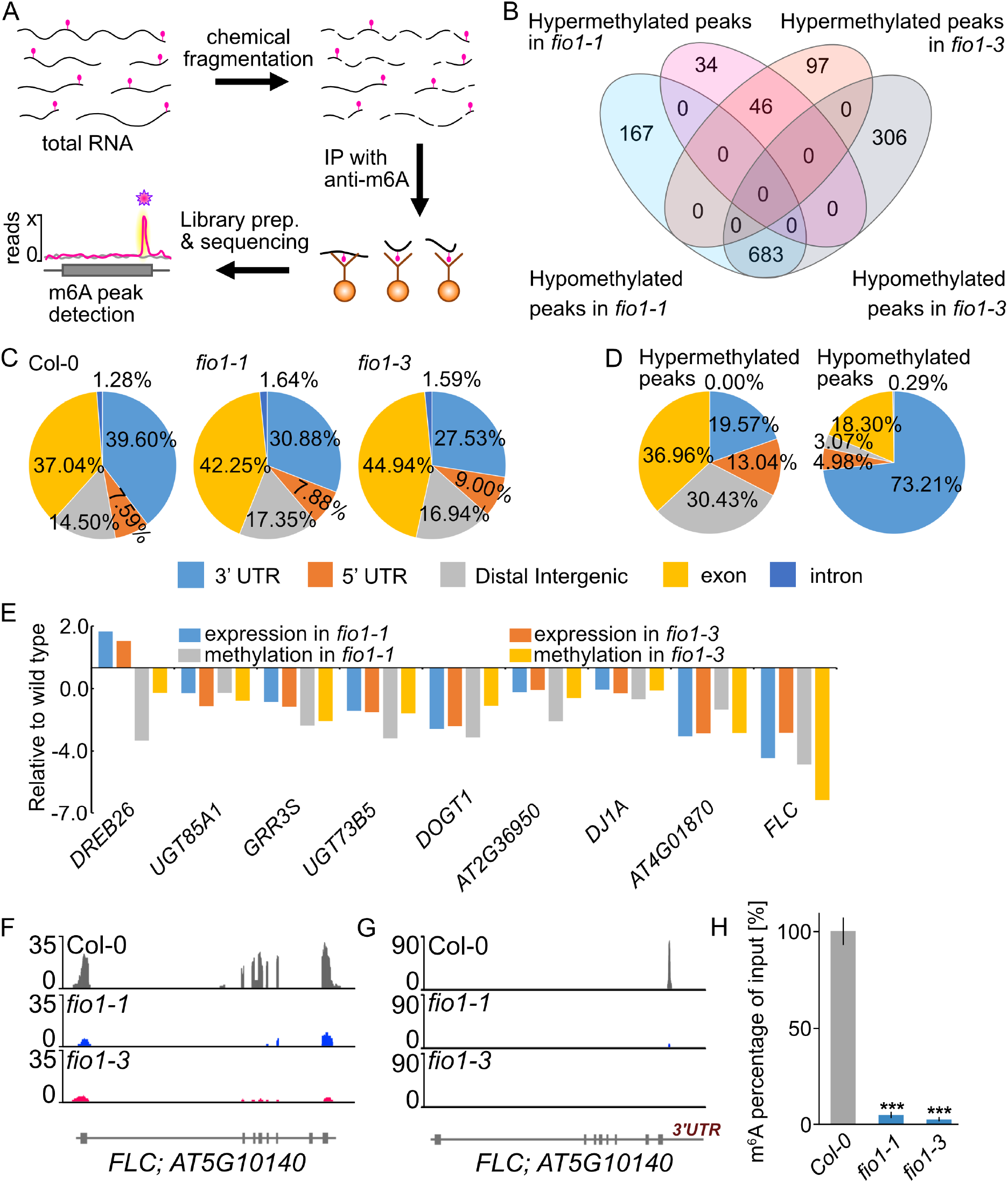
FIONA1 acts as m^6^A-methyltransferase in Arabidopsis. **(A)** Depiction of the meRIP-seq method. In brief, total RNA was isolated from seedlings and subsequently fragmented into small (100bp) fragments. After immunoprecipitation with an m^6^A-specific antibody, Illumina short-read sequencing libraries were generated and sequenced. After mapping all reads to the Arabidopsis genome, m^6^A peak regions (pink star) could be identified. **(B)** Venn diagram showing the overlap of the hypermethylated and hypomethylated m^6^A peaks identified in *fio1-1, fio1-3* compared to Col-0 wild type plants. **(C)** Comparison of distribution of m^6^A peaks in different segments of wild-type (left panel), *fio1-1* (middle panel) and *fio1-3* (right panel) transcripts. The panels show pie charts presenting the percentages of m^6^A peaks in different transcript segments. **(D)** Comparison of distribution of m^6^A peaks in different segments of differently methylated peaks (left panel), hypermethylated peaks (middle panel) and hypomethylated peaks (right panel) in the overlap of *fio1-1* and *fio1-3* compared to wild type. The panels show pie charts presenting the percentages of m^6^A peaks in different transcript segments. **(E)** Expression levels and m^6^A methylation levels of the transcripts in the overlapping of RNAseq and MeRIPseq. Gene expression levels were derived from RNA-Seq data. m^6^A methylation levels were derived from MeRIPseq data. **(F)** RNA-seq coverage observed at the *FLC* locus. RNA-seq reads in Col-0 (grey), *fio1-1* (blue) and *fio1-3* (pink). Gene model depicts exons and introns. **(G)** MeRIP-seq coverage observed at the *FLC* locus. RNA-seq reads in Col-0 (grey), *fio1-1* (blue) and *fio1-3* (pink). Gene model depicts exons and introns. **(H)** Percentages of the m^6^A methylated FLC mRNA in input samples in the wild type, *fio1-1* and *fio1-3* measured by m^6^A-IP-qRT PCR. Values are the means ±SD. N = 4, ***P≤ 0.001.

### FLC is a prime target of FIO1

The mRNA of the flowering repressor *FLOWERING LOCUS C* (*FLC*) was identified as a prime methylation target of FIO1 (Fig. 5e). We detected strongly decreased expression of *FLC* mRNA in *fio1* mutants compared to wild type (Fig. 5f) and the m6A peak that can be detected in wild type plants is absent in *fio1-3* and strongly reduced in *fio1-1* mutant plants (Fig. 5g). To verify that *FLC* is indeed a *bona fide* methylation target of FIO1, we performed anti-m^6^A antibody immunoprecipitations (m^6^A-IP) of total RNA from wild type (Col-0), *fio1-1* and *fio1-3* seedlings followed by qPCR (m^6^A-IP-qPCR). We found the relative amount of m^6^A methylated *FLC* mRNA was strongly decreased in both *fio1* mutant plants (Fig. 5h) confirming that FIO1 is the essential m^6^A methylase that methylates the 3’UTR of *FLC*.

### Direct RNA sequencing

To determine the genome-wide m^6^A methylation changes in *fio1* loss of function mutants compared to wild type and to validate *FLC* methylation and stability in an unbiased fashion, we employed Nanopore direct RNA sequencing. In Col-0 wild type plants, the majority (34.7%) of m^6^A methylations occurred in the GGACA element, followed by AGACT (27.2%), GGACT (22.9%) and GGACC (15.25) (Fig. 6A). In summary, our work defined the Arabidopsis consensus m^6^A methylation site as RGACH, in which R represents A or G and H all nucleotides except G, which corresponds with the RRACH element that had previously been identified (Luo *et al*., 2014). FIONA1 is a methylase that adds methyl-groups to adenine bases of RNAs. Messenger-RNAs that are targets of FIO1 are therefore expected to be hypomethylated in a situation of lost or reduced FIO1 activity. Our direct RNA-sequencing approach yielded 74 genes that were hypomethylated in *fio1-1* mutants compared to wild type and 63 genes in *fio1-3* (Fig. 6C and Supplementary Table 4). Another recent direct RNA-sequencing study of the fio1-2 knock-down mutant revealed over 2000 hypomethylated transcripts in Arabidopsis (Xu *et al*., 2022). The comparison with our datasets identified in total 28 hypomethylated transcripts that are detected in at least two mutants (Fig. 6C and Table 1).

**Figure 6.**
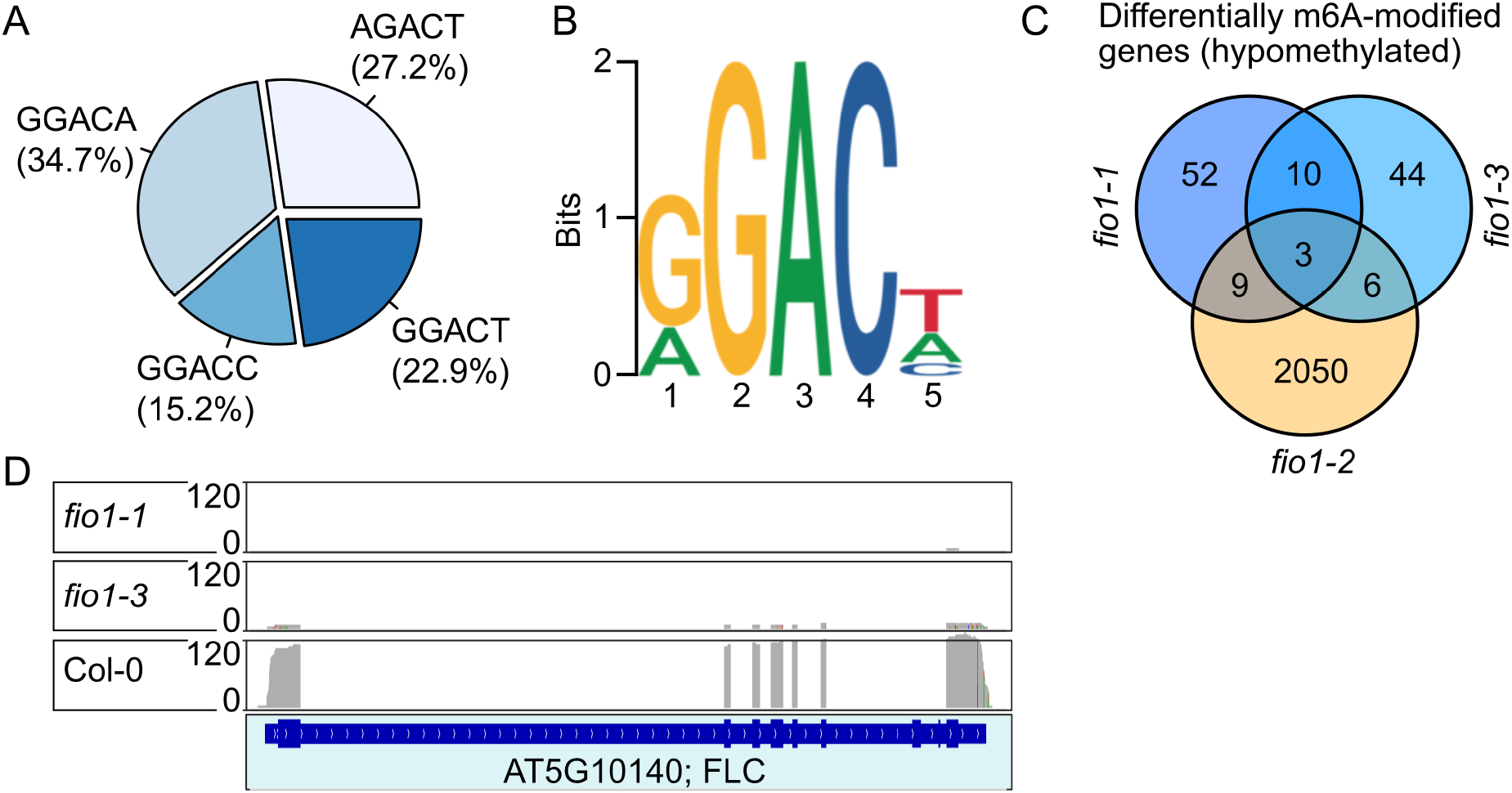
Direct RNA-sequencing analysis. **(A)** Distribution of m6A methylations detected by direct RNA-sequencing. **(B)** Logo of the conserved m6A sequence motif detected by direct RNA-sequencing. **(C)** Venn diagram showing the overlap of the hypomethylated m^6^A transcripts identified in *fio1-1, fio1-2* and *fio1-3* compared to Col-0 wild type plants. **(D)** Sequence coverage observed at the *FLC* locus. Direct RNA-seq reads in Col-0, *fio1-1* and *fio1-3*. Gene model depicts exons and introns.

**Table 1.** Comparative analysis of hypomethylated transcripts in *fio1-1, fio1-2* and *fio1-3* relative to wild type Col-0.

| Arabidopsis Gene Identifier (AGI) | Hypomethylated |  |  | Annotation |
| --- | --- | --- | --- | --- |
| AT1G12840 | *fio1-1* | *fio1-3* |  | DET3, ATVHA-C, ARABIDOPSIS THALIANA VACUOLAR ATP SYNTHASE SUBUNIT C, DE-ETIOLATED 3 |
| AT1G19980 | *fio1-1* | *fio1-3* |  | no symbol available |
| AT1G52040 | *fio1-1* | *fio1-3* |  | MBP1, ATMBP, myrosinase-binding protein 1 |
| AT1G52710 | *fio1-1* | *fio1-3* |  | no symbol available |
| AT1G76730 | *fio1-1* | *fio1-3* |  | COG0212, Clusters of Orthologous group 212 |
| AT2G18050 | *fio1-1* | *fio1-3* |  | HIS1-3, histone H1-3 |
| AT2G40480 | *fio1-1* | *fio1-3* |  | no symbol available |
| AT5G18790 | *fio1-1* | *fio1-3* |  | no symbol available |
| AT5G56860 | *fio1-1* | *fio1-3* |  | GNC, GATA21, GATA TRANSCRIPTION FACTOR 21 |
| AT5G64860 | *fio1-1* | *fio1-3* |  | AtDPE1, DPE1, disproportionating enzyme |
| AT1G50250 | *fio1-1* | *fio1-2* |  | FTSH1, FTSH protease 1 |
| AT1G52400 | *fio1-1* | *fio1-2* |  | BGL1, ATBG1, BGLU18, A. THALIANA BETA-GLUCOSIDASE 1 |
| AT1G63770 | *fio1-1* | *fio1-2* |  | no symbol available |
| AT2G30520 | *fio1-1* | *fio1-2* |  | RPT2, ROOT PHOTOTROPISM 2 |
| AT2G47940 | *fio1-1* | *fio1-2* |  | DEG2, DEGP2, EMB3117 \| DEGP protease 2, |
| AT3G10060 | *fio1-1* | *fio1-2* |  | no symbol available |
| AT3G51950 | *fio1-1* | *fio1-2* |  | no symbol available |
| AT5G42650 | *fio1-1* | *fio1-2* |  | CYP74A, AOS, DDE2, allene oxide synthase, DELAYED DEHISCENCE 2, CYTOCHROME P450 74A |
| AT5G66190 | *fio1-1* | *fio1-2* |  | LFNR1, ATLFNR1, FNR1, leaf-type chloroplast-targeted FNR 1, LEAF FNR 1 |
| AT1G67480 | *fio1-2* | *fio1-3* |  | no symbol available |
| AT2G22990 | *fio1-2* | *fio1-3* |  | SNG1, SCPL8, sinapoylglucose 1 |
| AT4G19110 | *fio1-2* | *fio1-3* |  | no symbol available |
| AT4G19160 | *fio1-2* | *fio1-3* |  | no symbol available |
| AT5G25265 | *fio1-2* | *fio1-3* |  | HPAT1, hydroxyproline O-arabinosyltransferase 1 |
| AT5G57560 | *fio1-2* | *fio1-3* |  | XTH22, TCH4, Touch 4, xyloglucan endotransglucosylase/hydrolase 22 |
| AT2G01490 | *fio1-1* | *fio1-2* | *fio1-3* | PAHX \| phytanoyl-CoA 2-hydroxylase |
| AT2G28900 | *fio1-1* | *fio1-2* | *fio1-3* | OEP16, OEP16-1, ATOEP16-L, ATOEP16-1, outer plastid envelope protein 16-1 |
| AT4G08950 | *fio1-1* | *fio1-2* | *fio1-3* | EXO, EXORDIUM |

*FLC* expression was shown to be significantly reduced in both *fio1-1* and *fio1-3* mutants and meRIP-seq detected m^6^A methylation in the 3’UTR of *FLC* (Fig. 4F and Fig. 5F,G). In agreement with these latter results, direct RNA-sequencing confirmed that *FLC* mRNA is depleted in both *fio1-1* and *fio1-3* mutants (Fig. 6D).

## DISCUSSION

The precise timing of the floral transition is crucial for reproductive success. Premature as well as delayed flowering can result in seed dispersal at times where the offspring will be facing suboptimal conditions for survival and reproduction. This could either be due to the absence of pollinators or adverse environmental conditions. Therefore, a highly integrative network of transcription factors, but also epigenetic regulators, operate to ensure that flowering occurs in the most optimal conditions.

Methylation of mRNA is crucial for various functions within the cell. The m^6^A methylation of mRNA is an ancient molecular process and its disruption strongly compromises cellular functions. Strong reduction of the global m^6^A methylome early in plant development, as seen in mutants lacking the *METTL3*-homolog *MTA*, causes embryonic arrest (Zhong *et al*., 2008). Partial complementation of the *mta* mutant resulted in plants with compromised m^6^A levels that showed pleiotropic phenotypes such as reduced apical dominance and missing floral organs (Bodi *et al*, 2012). These latter results suggest that more subtle reductions of the global m^6^A levels are not detrimental to plant development. We provide further support of this by showing that the loss-of-function mutants of FIO1, a protein that is not essential for plant development, have only a subtle effect on the global m^6^A-methylome.

Furthermore, in contrast to the effect the loss of its homolog has on animal development, FIO1 is not essential and causes hypomethylation of specific transcripts. These hypomethylated mRNAs can then be stabilized, or destabilized, or mis-spliced. Affected transcripts encoding transcription factors or other regulators can subsequently induce alterations of circadian rhythms, cause changes in the production of hormones, or misregulation of other biological processes. Consistent with these multifaceted changes is the pleiotropic phenotype of *fio1* mutant plants. The precocious flowering phenotype is the most striking but *fio1* mutants additionally display a constitutive shade-avoidance phenotype, earlier senescence, and paler leaves (Kim *et al*., 2008). In accordance with these phenotypes, our RNA-seq study revealed that several genes encoding circadian clock regulators and positive regulators of flowering time were upregulated in the *fio1* mutant background (e.g. *LHY, PIF4*). In contrast, several of the downregulated transcripts encoded transcription factors that repress flowering (Fig. 1D, Supplementary Table 1).

Genetically, flowering is controlled by distinct pathways that interact at multiple levels to integrate inputs from all pathways. This integration ensures flowering occurs at the optimal time. The photoperiod pathway controls flowering in response to daylength and involves the B-Box zinc finger transcription factor CONSTANS (CO) which, in Arabidopsis, is stabilized at the end of long days (Valverde *et al*., 2004). CO positively regulates the expression of *FLOWERING LOCUS T (FT*) (Samach *et al*, 2000), encoding a mobile protein that travels to the shoot meristem to induce flowering (Corbesier *et al*., 2007). *FIO1* acts partially through the photoperiod pathway and the early flowering phenotype of *fio1* mutants correlates with increased levels of both *CO* and *FT* mRNAs (Fig. 3A,B) as well as increased levels of *SUPPRESSOR OF OVEREXPRESSION OF CONSTANS1* (*SOC1*) (Xu *et al*., 2022). Our genetic interaction studies have shown that mutations in both *CO* and *FT* can partially suppress the early flowering effect of *fio1* mutants. Consistent with our findings, the *soc1* mutant has also been shown to partially suppress the early flowering phenotype of *fio1-2* mutant plants (Xu *et al*., 2022). Taken together, these data support a model that assumes an indirect effect of the photoperiod pathway in the control of flowering by FIO1.

Our RNA-sequencing data identified both up- and downregulated transcripts in *fio1* mutants compared to wild type. However, the overlap between the set of de-regulated transcripts identified in *fio1-2* mutants (Xu *et al*., 2022) is very limited. The latter fact can be attributed to the different types of mutations that were analyzed. While our study capitalized on loss-of-function mutants, *fio1-2* is a T-DNA insertion line that still expresses *FIO1* mRNA, although at a lower level. Alternatively, the observed differences could be technical in nature, the result of either of the different sequencing approaches that were chosen or the growth conditions in which plants were cultivated.

MeRIP-sequencing further confirmed that FIO1 is likely not the main factor in the m^6^A modification of mRNAs but a more selective methylase that modifies specific mRNAs. This assumption is supported by the finding that loss-of-function mutants are viable and able to produce fertile offspring. Interestingly, despite the much higher number of differentially methylated transcripts in the *fio1-2* mutant (Xu *et al*., 2022), the comparison of the differentially hypomethylated transcripts compared to those in *fio1-1* and *fio1-3* (this study) produced only a very moderate overlap (Fig. 6C). Again, this might be due to the application of different methods or an indication that the reduction of FIO1 activity affects the m^6^A methylome more strongly than does the complete loss. Furthermore, the analysis of the m^6^A consensus in *fio1-2* identified the YHAGA motif, which is significantly different to the RRACH motif that has been described in both plants and animals (Luo *et al*., 2014; Warda *et al*, 2017), and to the RGACH consensus sequence that we identify in this work(Fig. 6B).

Detailed analysis of specific transcripts that are differentially methylated and differentially expressed identified the flowering regulator *FLC*. Regardless of whether the contribution of *FLC* methylation contributes only marginally to the early flowering response of *fio1* mutants, our work unequivocally demonstrates that FIO1 is the m^6^A-methylase that methylates the 3’UTR of *FLC* mRNA. We speculate that the failure to methylate *FLC* mRNA targets it for rapid degradation, hence the absence of *FLC* mRNA in *fio1* mutants. In any case, further characterization of the relationship between FIO1 and the biology of *FLC* will lead to insights into the function of its 3’-end methylation.

Our analyses focused on the role of methylation of mRNAs and the impact on the regulation of flowering. We cannot rule out confounding effects that the loss of *FIO1* may have on the methylation and regulation of the non-coding transcriptome. Such effects might also contribute to the phenotype of *fio1* mutant plants and further characterization is needed to shed light on these processes.

## METHODS

### Plant materials and growth conditions

*Arabidopsis thaliana* genotypes used in the study were, if not otherwise stated, in the Columbia Col-0 background. Double and triple mutant plants, such as *fio1 co-sail, fio1 ft10* and *fio1 miP1a miP1b* were generated by genetic crossing. For flowering experiments, seeds were stratified 48 h at 4°C, and grown on soil in a plant growth chamber under long daylight conditions (16 h light / 8 h dark), or short daylight conditions (8 h light / 16 h dark) at 22 °C day / 20 °C night. Flowering time was measured by counting the number of rosette leaves at the bolting stage.

For RNA-seq, MeRIP-seq and qPCR, 14-day old seedlings were collected. Seeds were sterilized in 70% ethanol and sown on 1/2 Murashige and Skoog (MS) medium plates with 0.8% agar and kept at 4°C for 48 hours in darkness for stratification and then grown at (22 °C day / 20 °C night) and 70% humidity under long daylight conditions (16 h light / 8 h dark).

Loss-of-function mutants of *fio1* were generated using the CRISPR/Cas9 vector pKI1.1R, containing the Cas9 expression cassette (RPS5Ap::Cas9:HspT), a sgRNA expression cassette (U6.26p::AarI_site:sgRNA) and, for selection the RFP expression cassette (OLE1p::OLE1:TagRFP). Single-guide RNAs (sgRNAs) were designed using the web tool CRISPR-P v 2.0 (Liu *et al*, 2017). Vectors with sgRNAs were generated according to the published description (Tsutsui & Higashiyama, 2017). To create mutants with deletions, two to three Agrobacterium strains GV3101 pMD90 with different sgRNAs (Supplementary Table. 3) were pooled and transformed into wild type plants via floral dip. RFP-positive seeds were selected using a Leica MZFLIII stereomicroscope equipped with RFP filters. Deletions were detected by PCR based sequencing.

### Mapping-by-sequencing

91.99% sequenced reads were mapped by Bowtie2 (v2.1.0)(Langmead & Salzberg, 2012) using the TAIR9 genome assembly and TAIR10 annotation from Phytozome v10.3 (phytozome.org). SNP calling was performed using samtools and BCFtools (v0.1.19)(Li, 2011; Li *et al*, 2009). 1118 (Chr1: 203, Chr2: 194, Chr3: 247, Chr4: 189, Chr5: 285) background corrected EMS-induced SNP markers were identified by SHOREmap(Schneeberger *et al*, 2009) (v3.2) using standard settings. Finally, the mutations indicated a mapping interval of 7 Mb Kb on chromosome 2, containing 84 mutations. The trend line is the average of all SNP allele frequencies in a sliding window (size: 2,500 Kb; step: 100 Kb).

### FIO1 homology modeling

The methyltransferase domain of FIONA1 (UniProt accession code F4IGH3, residues 1-333) was modelled with Phyre2 (http://www.sbg.bio.ic.ac.uk/phyre2) using the Intensive modelling mode. The resulting homology model was aligned against the human crystal structure of the human FIONA1 homologue, METTL16 (PDB ID: 6DU4) for structural analysis.”

### mRNA sequencing analysis

For RNAseq analysis, we collected two biological replicates of 14 day-old wild type (Col-0), *fio1-1, fio1-3* seedlings. Total RNA was extracted from 100 *A. thaliana* seedlings for each line grown on a ½ MS agar plate using the SpectrumTM Plant Total RNA Kit (Sigma-Aldrich) following the manufacturer’s instructions. Total RNA was treated with DNAase I (RapidOut DNA Removal Kit, Thermo Scientific) according to the manufacturer’s instructions. Sequencing library preparation and sequencing on an Illumina HiSeq4000 instrument was performed by Novogene (Hongkong). About 3.7 Gb high-quality 150-bp paired-end reads were generated from each library. FastQC (Galaxy Version 0.72 + galaxy1) was initially run to assess the overall quality of all sample reads. Poor quality bases and adapters were filtered out using Trim Galore (Galaxy Version 0.6.3). The quality-filtered reads were aligned to the *Arabidopsis thaliana* reference genome (TAIR10) using HISAT282 (Version 2.1.0 + Galaxy4) with default parameters. HTseq (Galaxy Version 0.9.1) software was used to count the number of raw reads mapped to each of the genes. Differential expression analysis was performed with four analytical methods, DEseq 2 (Galaxy Version 2.11.40.6+galaxy1), edgeR (Galaxy Version 3.24.1+galaxy1), Limma-voom (Galaxy Version 3.38.3+galaxy3) and Limma-trend (Galaxy Version 3.38.3+galaxy3). All four statistical methods gave similar overall conclusions. We selected the most conservative results (Limma-voom; false discovery rate (FDR) = 0.05) for further investigation. Significance testing was performed using the Benjamini-Hochberg method(Benjamini & Hochberg, 1995). Genes showing an absolute value of log2 FC (fold change; *fio1* mutant / WT) ≥ 1.0 and adjusted P-value (false discovery rate; FDR) < 0.05 were considered as differentially expressed genes. RNAseq data generated in this study has been deposited in NCBI’s Gene Expression Omnibus under GEO Series accession no. GSE171926.

### m6A RNA Immunoprecipitation sequencing (MeRIP-seq) and data analysis

MeRIP-seq was performed as described before(Dominissini *et al*., 2013) with modifications. Briefly, total RNA was extracted from 14 day-old Arabidopsis thaliana seedlings using the SpectrumTM Plant Total RNA Kit (Sigma-Aldrich) and treated with DNAase I (RapidOut DNA Removal Kit, Thermo Scientific). 300 μg of total RNA was mixed with 10×Fragmentation buffer (1 M Tris-HCl pH=7.0, 1 M ZnCl2) and placed at 94 °C for 5 min then snap cooled on ice for 5 minutes. The volume of fragmented RNA was then adjusted to 755 μl with RNase-free water. Next, 10 μL RNasin Plus RNase inhibitor (Promega, cat. no. N2611), 10 μL Ribonucleoside vanadyl complexes (RVC; 200 mM; Sigma-Aldrich, cat. no. R3380), 200 μL 5×IP buffer (50 mM Tris-HCl, 750 mM NaCl and 0.5% (vol/vol) Igepal CA-630), and 25 μL of m6A antibody (Synaptic Systems, cat. no. 202 003) were added to samples and samples were rotated at 4°C for 2 hours. After 2 hours, pre-blocked Protein A Dynabeads™ (Thermo Fisher, 1001D) was added to the RNA samples and rotated for an additional 2 hours at 4°C. After 2 hours, Dynabeads were pelleted using a magnetic stand and washed three times with 1 mL 1×IP buffer. RNA was eluted from Dynabeads by adding 98 μL elution buffer (20 mM Tris-HCl pH 7.5, 300 mM sodium acetate, 2 mM EDTA, 0.25% SDS), 2 μL of proteinase K (Thermo Fisher, AM2546) and then shaking for 1 hour at 37°C. All samples were precipitated using 3 M sodium acetate (pH 5.2) and 2.5 volumes of 100% ethanol and kept at −80°C overnight. Libraries were prepared using NEBNext Multiplex Small RNA Library Prep Set for Illumina (New England BioLabs, E7300S) according to the manufacturer’s instructions. Novogene (Beijing) performed sequencing on an Illumina HiSeq4000 instrument. About 3.0 Gb high-quality 150-bp paired-end reads were generated from each library. FastQC (Galaxy Version 0.72 + galaxy1) was initially run to assess the overall quality of all sample reads. Poor quality bases and adapters were filtered out using Trim Galore (Galaxy Version 0.6.3). The quality-filtered reads were aligned to the *A. thaliana* reference genome using HISAT2 (Version 2.1.0 + Galaxy4) with default parameters. To identify regions in which m6A modifications occurred, MACS (Zhang *et al*., 2008) was used to call peaks on aligned files. The peaks showing an absolute value of log2 FC (fold change; *fio1* mutant / WT) ≥ 1.0 and raw reads ≥ 50 were considered as differentially modified peaks. MeRIPseq data generated in this study has been deposited in NCBI’s Gene Expression Omnibus under GEO Series accession no. GSE171928.

### Nanopore direct RNA sequencing

Total RNA was isolated as described above for mRNA-seq and direct RNA sequencing libraries were prepared by CD genomics using the Oxford Nanopore DRS protocol (SQK-RNA002, Oxford Nanopore Technologies). Samples were loaded into the Nanopore R9.4 sequencing micro-array and sequenced for 48-72 hrs using the PromethION sequencer (Oxford Nanopore Technologies). Read quality assessment, base calling and adapter trimming was carried out with the Guppy software (version 3.2.6). Nanofilt (version 2.7.1) was then used to remove low quality reads (Q-value < 7) and short-length reads (<50 bp). The clean reads were subsequently corrected using Fclmr2 (version 0.1.2). Minimap2 (version 2.17-r941) was used to map the clean reads to the *A. thaliana* genome and the alignment ratio of clean reads to the reference genes was calculated using Samtools (version 1.10). To identify m6A sites, the Tombo software de novo model together with MINES was used for calculation. Methylkit software was then used to analyze differential methylation sites (DML). Logistic regression test was used to detect differential methylation sites.

### RNA m^6^A immunoprecipitation RT-qPCR

Quantitative real-time PCR was performed to assess relative abundance of m6A RNA in the RIP samples. 300 μg total RNA was adjusted the volume to 1000 μl with 5×IP buffer (50 mM Tris-HCl, 750 mM NaCl and 0.5% (vol/vol) Igepal CA-630) and RNase-free water and incubated with 10 μg m6A antibody (Synaptic Systems, cat. no. 202 003, Goettingen, Germany). The mixture was rotated at 4 °C for 2 h, then pre-blocked and washed Dynabeads™ Protein A (Thermo Fisher, 1001D) were added and the mixture rotated for an additional 2 h at 4 °C. After washing with IP buffer containing Ribonucleoside vanadyl complexes (RVC, Sigma, R3380-5ML) three times, the m6A IP RNA was eluted with 98 μL elution buffer (20 mM Tris-HCl pH 7.5, 300 mM sodium acetate, 2 mM EDTA, 0.25% SDS). 2 μL of proteinase K (Thermo Fisher, AM2546) was added and the RNA incubated for 1 hour at 37°C with gentle shaking. All samples were precipitated using 3 M sodium acetate (pH 5.2) and 2.5 volumes of 100% ethanol and kept at −80°C overnight. cDNA was synthesized by iScript™ cDNA Synthesis Kit (Bio-Rad). qPCR analyses was done with Ultra SYBR Mixture with ROX (CWBIO) on a CFX384 Touch Real-Time PCR Detection System (Bio-Rad). qRT-PCR primers that were used to amplify *FLC* were: flc_qF: AGCCAAGAAGACCGAACTCA and flc_qR: TTTGTCCAGCAGGTGACATC.

## DATA AVAILABILITY

All data has been submitted to public repositories and the respective links have been included in the respective sections of the material and methods.

## ACKNOWLEDGEMENTS

We acknowledge funding through NovoCrops Centre (Novo Nordisk Foundation project number 2019OC53580 to S. W.), the Independent Research Fund Denmark (0136-00015B and 0135-00014B to S. W.), the Novo Nordisk Foundation (NNF18OC0034226 and NNF20OC0061440 to S. W.).

## AUTHOR CONTRIBUTIONS

BS, KKB and SW designed the study; BS, KKB, AE, LP, VK, AB, UD, VR and DS performed experiments; BS, KKB, DS and SW analyzed the data; SW provided supervision and wrote the manuscript with input from all co-authors.

## SUPPLEMENTARY MATERIAL

### SUPPLEMENTARY FIGURES

**Figure S1.**
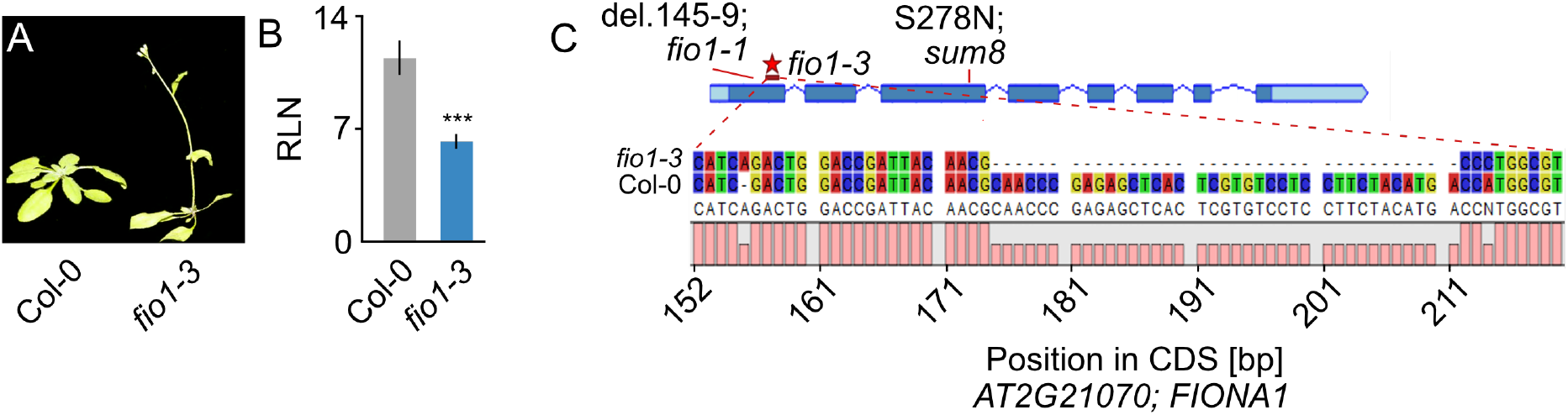
Analysis of *fio1-3*, a CRISPR-induced mutation in *FIO1*. **(A)** Phenotype of *fio1-3* compared to the Col-0 wildtype when grown in LD conditions. **(B)** Determination of flowering by counting the number of rosette leaves (RLN = rosette leaf number) at the bolting stage in LD. Plotted are average leaf number +/- SD, ***p=<0.001, N=10-14. **(C)** Nucleotide alignment showing the CRISPR-induced genomic deletion found in *fio1-3*. Gene model on top shows the relative positions of all three *fio1* mutations.

**Figure S2.**
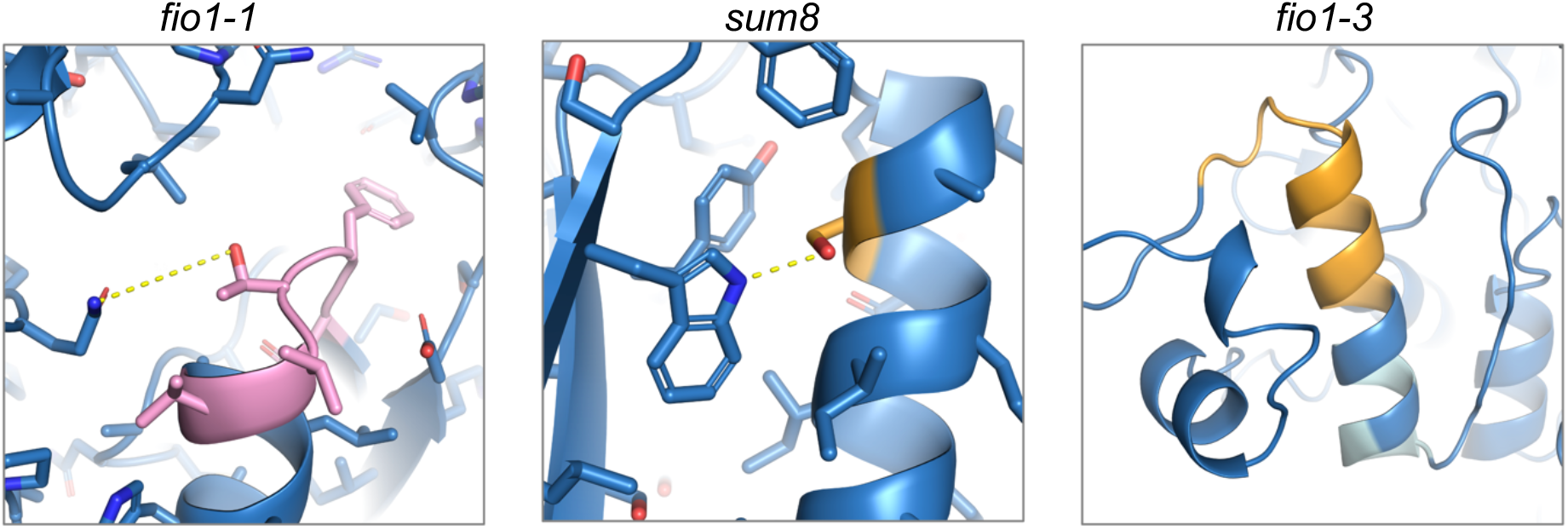
Analysis of FIO1 methyltransferase domain mutants based on homology model. The three mutants were mapped to the homology model of FIO1 (see Materials and Methods). The *fio1-1* mutation involved the loss of five amino acids highlighted in pink, including the loss of a potential hydrogen bond between the threonine and asparagine. The *sum8* mutation changes the serine (orange), which normally hydrogen bonds to a tryptophan, into an asparagine. The resulting larger side-chain of asparagine is unlikely to be accommodated in the constrained protein interior, leading to changes in the protein structure and loss of function. The *fio1-3* mutation involves a large deletion (orange) and missense mutations (light cyan) in a partially buried alpha helix, which are very likely to disrupt protein folding and function.

**Figure S3.**
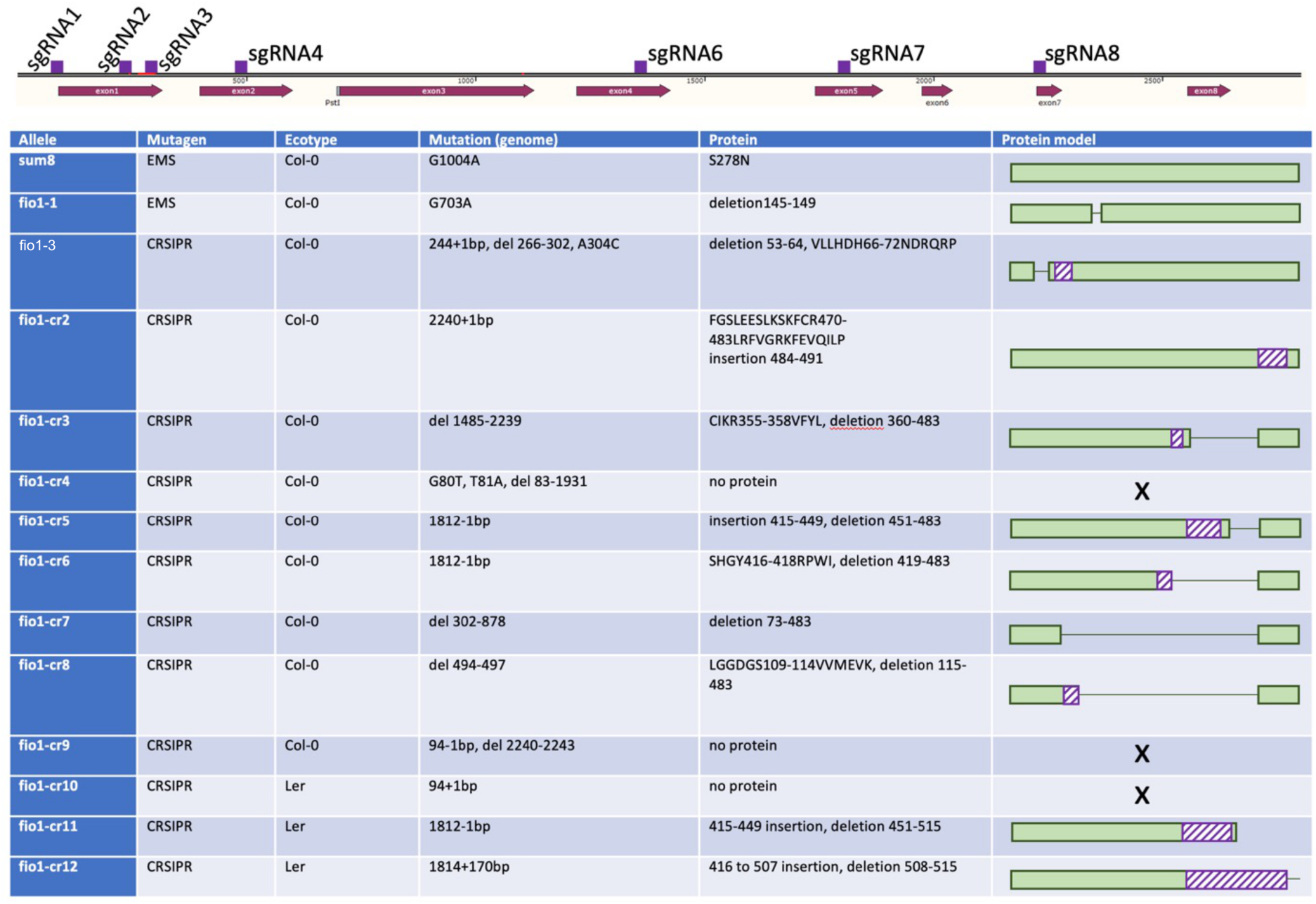
Overview of additional CRISPR-induced mutations in *FIO1*. Gene model depicting the *FIO1* locus (exons in dark read and location of sgRNAs in purple). All sgRNAs were transformed in bulk and from all early flowering individuals the *FIO1* gene was sequenced to determine the nature of CRISPR-induced mutations. To determine the correct reading frame, RNA was isolated and *FIO1* was amplified on cDNA and subsequently sequenced.

### SUPPLEMENTARY TABLES

**Supplementary table 1: DEGs identified in *fio1-1* and *fio1-3* by RNAseq.**

**Supplementary table 2: Methylation peakes identified by MeRIP-seq.**

**Supplementary table 3: Oligonucleotide sequences.**

**Supplementary table 4: Hypomethylated transcripts identified in *fio1-1* and *fio1-3***

